# Epidemiological Study Of Foot And Mouth Diseases Through Serological And Molecular Investigation In Cattle Of Selected Districts, Jimma Zone, Southwest Ethiopia

**DOI:** 10.1101/2024.06.11.598420

**Authors:** Hailehizeb Tegegne, Seid Ababulgu, Eyoel Ejigu

## Abstract

Foot and mouth disease is highly contagious and notifiable transboundary disease of cattle that can cause a huge cattle productivity and production loss. A cross-sectional study was performed to estimate sero prevalence, assess associated risk factors and molecular detection of FMDV in cattle. Cluster sampling technique was employed for the selection of sampling units for the seroprevalence study. A total of 245 blood samples were collected using plain vacutainer tubes and the obtained sera were tested by 3ABC-Ab ELISA at the Animal Health institute. Twenty nine (29) epithelial tissue and vesicular fluid samples were collected purposively from outbreak cases for the molecular detection of FMDV. Kebeles and individual cattle were randomly selected, while households were designated using systematic random sampling method. An overall prevalence of 22.45% (95%, CI=17.22%-27.67%) was recorded. Multivariable logistic regression analysis indicated that herd size, age, new animal introduction into the herd and management system were the major risk factors, significantly associated with FMD sero positivity (P<0.05). The large herd size had 4-times (OR=3.97; P=0.000) more odds of FMD sero-positivity compared to the small herd sizes. The FMD seropositivity decreases 0.11013 as the Cattle age increases by 5years with the (coefficient=-0.11; P=0.172). The animals from herd to which new animals was introduced had nearly 9-times more odds (OR=9.40; P=0.000) of sero positivity than the animals sampled from no new animal introduction. Likewise, cattle those reared under extensive management system were 4-times (OR=4.10; P=0.009) at higher chance of being sero-positive compared to the intensive one. From outbreak cases, 27 (93.1%) were identified positive for FMDV serotype SAT 2. A total of 124 individuals were interviewed, and the majority responded that there is no practice of reporting disease outbreak, free animal movement, free rangeland grazing and they use traditional case management as a means of controlling the disease. The finding of FMD virus antibodies in cattle from all study areas indicate endemic circulation of the virus. The implementation of regular vaccination could minimize the occurrence and further molecular characterization should be needed to identify other serotypes of FMD virus that could inform to supply an appropriate vaccine to the area.

## 1. INTRODUCTION

Ethiopia has approximately 59.5 million cattle, 30.7 million sheep and 30.2 million goats (CSA, 2021). The contribution of the livestock sector to the national economy is minimal compared to its potential. One of the main reasons for this is the widespread occurrence of many infectious diseases, such as foot and mouth disease (FMD), which drastically reduces the production and productivity of livestock ((CSA, 2021).

Foot and Mouth Disease is the most widespread and highly contagious transboundary viral disease of cloven-hooved animals. It imposes significant economic impact, in cattle and swine as well as sheep and goats (Mohaptra *et al*., 2018: Ali *et al*., 2019). It is the most important disease of livestock because of its severe impact on production and trade of animal and animal products (Thomson, 2014). It was the first disease for which the OIE established official status recognition. It caused by Foot and mouth disease virus (FMDV), a virus under the genus Aphtho virus, family of Picorna viridae, a positive-sense single-stranded RNA virus with a very low molecular weight ranging from 7.2 to 8.4 kb (Moustafa, 2018). It has very simple and small structure, 25-30 nm diameter in size, which accelerates the air transmission of the virus, allowing it to spread over long distances in a very short time by following the nature of the wind speed and direction (Ashenafi *et al*., 2019).

An icosahedral capsid layer surrounded the viral genome that composed of four structural proteins, namely, VP1, VP2, VP3, and VP4 with 60 copies of each protein. FMDV has a high mutation rate because the viral RNA dependent RNA polymerase lacks proofreading ability. There are seven serotypes of FMD virus: A, O, C, Asia 1, and SAT 1-3 (South African territory) (Ali *et al*., 2019; OIE, 2017). All the serotypes produce a disease that is clinically indistinguishable but immunologically distinct. Currently, four of the seven serotypes of FMDV (O, A, SAT 1, SAT 2) are endemic in Ethiopia, while serotype C was last diagnosed in 1983 (Sulayeman *et al*., 2018).

As to the epidemiological eyeglass, and from disease control perspectives, each serotype causes immunological distinctive diseases. As there is no cross-immunity between serotypes, immunity to one type does not confer protection against the others, and protection from other strains within a serotype varies with their antigenic similarity (Anna *et al*., 2015; Lycett *et al*., 2019). Analysis of strains of FMD virus by antigenic and genetic profiles is important in epidemiological studies and for the selection of the most appropriate vaccine strains for a region where vaccination is practiced (FAO/OIE, 2018).

Foot and Mouth Disease is transmitted by a variety of methods between herds, countries and continents but, the main route of infection to susceptible animal is either orally (mainly for swine) or via the respiratory tract (mainly cattle). Aerosol transmission is the major means of animal-to-animal spread within premises. Vesicular lesions and erosions of the epithelium of the mouth, nose, muzzle, feet, teats and udder characterize the disease. FMD-infected animals usually develop blister-like lesions in the mouth, tongue and lips, teats, or between the hooves, which causes them to salivate profusely or become lame (Amass *et al*., 2004; Yalew, 2019). It has also, transboundary potential that is responsible for the outbreak. Livestock movements in inter-country are important factors for spreading the virus and the virus can travel as far as 250 km via aerosol (Abdula *et al*., 2011; Dee *et al*., 2018).

Foot and Mouth Disease widely distributed and endemic in Africa, Asia, South America and parts of Europe. The disease can occur in any country but Japan, New Zealand and Australia are disease free (FAO, 2004; Yalew, 2019). In Ethiopia FMDV is endemic in all production systems and a large number of outbreaks were reported every year (Ayelet, *et al.,* 2012; Jemberu *et al*., 2016). Majority of the outbreaks were caused by serotypes O, A, SAT 2 and SAT 1 (Jemberu *et al*., 2016; Yalew, 2019). Studies of FMD undertaken in different parts of the country so far revealed the existence of the disease and reported seroprevalence ranging from 5.6%-44.2% % (Jenbere *et al*., 2011)

Effective control and eradication of FMD mainly depends on accurately and timely diagnosis of the disease in endemic areas and in backing of stamping out policies in FMD free regions. For an efficient and rapid diagnosis of FMD, reverse transcriptase polymerase chain reaction is the most widely used techniques in parallel with conventional assays such as virus isolation, serology, and virus neutralization test (Xu *et al*., 2013; Seoke, 2016). The current measures to control FMD include vaccination using matching vaccine, movement control, and slaughter of infected or susceptible animals. However, antibody generated by infection or vaccination against one serotype fails to cross-protect against the other serotypes (Domingo *et al*., 2003). Consequently, vaccine strain requirements differ according to the types and subtypes of virus and vaccines have to be selected with great care (Mumford, 2007).

### Statements of the Problem

There were multiple FMD outbreaks in and around Jimma. However, there was no any study done concerning the disease prevalence at the molecular level. Different types of FMDV lineages or serotypes cause these outbreaks. It was likely that the control measures put in place following an outbreak in the study area were not adequate, so the disease spread to nearby areas through live animal movement since there was no legal framework put in action to restrict illegal animal trafficking in the area. Epidemiological investigation of FMD outbreaks can give insight about disease patterns, which might be used for early warning and prospective control planning of the disease. Delayed case detection enhances disease transmission and epidemic risk, and subsequent economic losses. Comprehensive epidemiological knowledge of FMD is crucial for the development of efficacious surveillance and control programs. Foot and Mouth Disease is endemic Jimma Zone. However, epidemiological knowledge related to FMD was scarce, due to inefficient surveillance resulting in under reporting of FMD disease. Extensive published data was not available about the molecular epidemiology and risk factors of FMD outbreaks in the study area and to achieve the goal of FMD control strategy, a well-established knowledge of disease determinants is required. Since molecular characterization of FMDV is not routine in the study area, the serotype and the genetic relationships among the viruses responsible for FMD outbreaks is not known. It is not known whether the outbreaks in the study sites are due to a common source or they are new and independent introductions. Due to the highly infectious nature of the virus and the accompanying economic constraints following, an outbreak of the disease, surveillance and characterization of the virus is crucial.

Along with this knowledge gap, there is no government nationally programmed strategy to control FMD through vaccination and/or movement control in the country. Lack of vaccination strategies (quality, coverage and timing) and presence of free animal movement without certification are thus the main factors that could increase the spread of FMD along the cattle market chain. Effective control strategies of this disease need sensitive, specific and rapid diagnostic tools like molecular detection (real time RT-PCR), virus isolation and antigen detection by ELISA. However, in Ethiopia the information and knowledge of molecular characterization and serotyping is scarce particularly in remote areas of the country. For the development of acceptable FMD control and prevention, proper outbreak investigation, identification and characterization of the circulating serotype of the virus, and assessing the associated risk factors of the outbreaks is required.

Therefore, the objectives of this study were:

- To identify sero prevalence of FMD and potential risk factors associated in selected districts of Jimma zone
- Molecular detection and identify the circulating serotype of the FMD virus from outbreak cases during the study.

## 2. MATERIALS AND METHODS

### 2.1. Description of the Study Area

The study was conducted in two areas of Jimma Zone, South West Ethiopia. Jimma zone is situated at a distance of 356 Km, South West of Addis Ababa. It lies between 7°41“N latitude and 36°50” E longitudes and at an average elevation 1750 masl above sea level. The climate of the Zone is characterized by humid tropical with bimodal heavy rainfall, which is uniform in amount and distribution, ranging from 1200 to 2800mm per year, with short and main seasons occurring from mid-February to May and June to September, respectively. The mean annual minimum and maximum temperature ranging from 6°C and 31°C respectively. The overall average temperature is approximately 18.5 °C. Jimma zone is one of the largest in livestock populations in Ethiopia with cattle population estimated 2,212,962 heads. Mixed crop livestock production is the main agricultural activity in the area (CSA, 2021).

Dedo is one of the woredas in the Oromia Region of Ethiopia. Part of the Jimma Zone, Dedo is bordered on the south by the Gojeb River and South Western Region of Ethiopia that recently formed and which separated from the Southern Nations, Nationalities and Peoples Region, on the west by Seka Cokorsa (woreda), on the north by Kersa, and on the east by Mencho (woreda) that recently separated from Dedo Woreda. It is located between 7°13’-7°39’ north latitudes and 36°43’-37°12’ east longitudes. The altitude of this woreda ranges from 880 to 2500 meters above sea level. Crop-livestock production is the major farming system practiced in the district.

Jimma city is located in southwestern Oromia Region, Ethiopia on 365 km distance from Addis Ababa. It is a special zone of the Oromia Region and is surrounded by Jimma Zone. It has a latitude and longitude of 7°40′N 36°50′E. It lies at an elevation of 5,740 feet (1,750 meters.a.s.l) in a forested region known for its coffee plantations. Jimma serves as the commercial center for the region, handling coffee and other products. An agricultural school and an airport serve the town.

**Figure 1:**
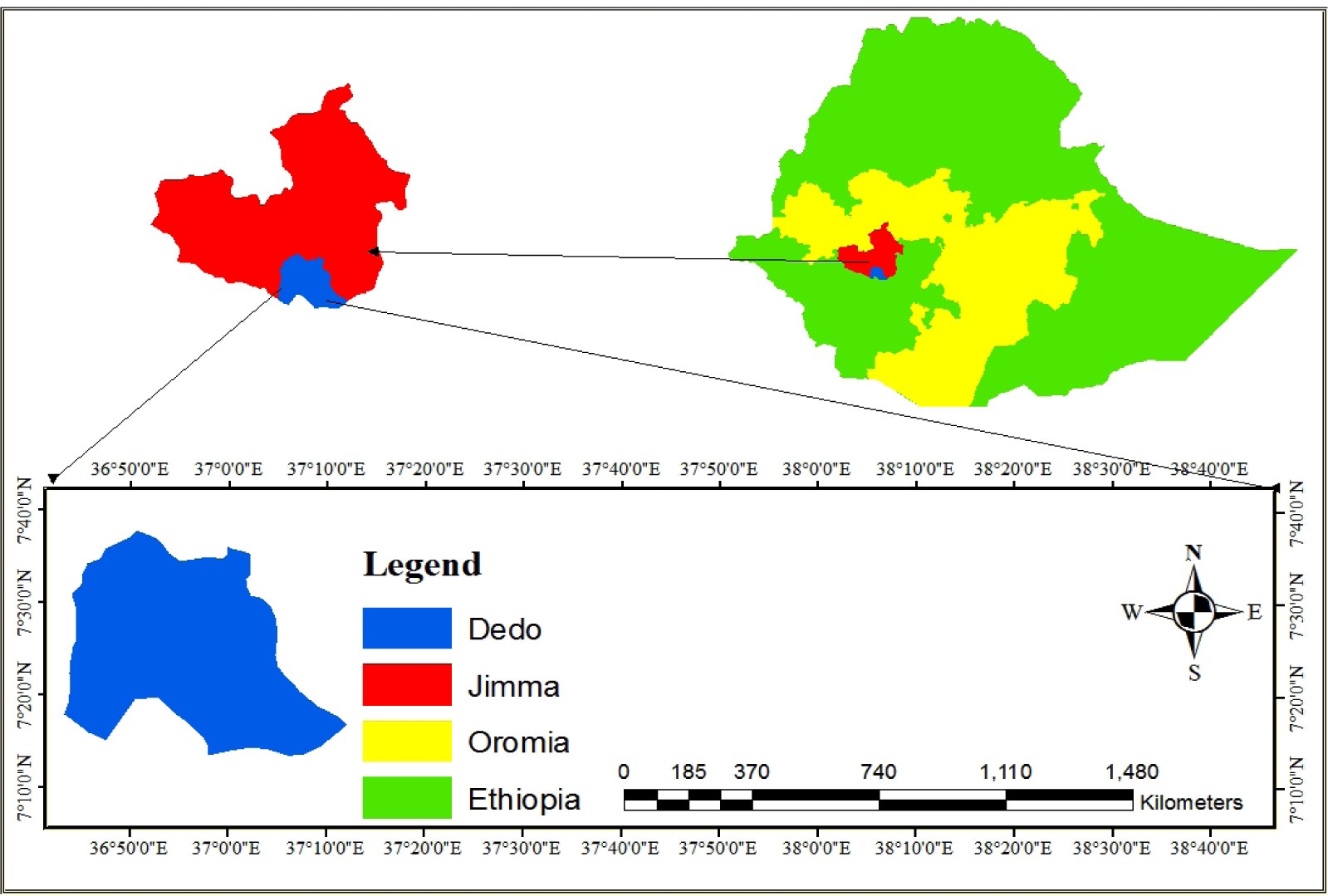
Map of the Study Area.

The study area was selected based on the presence of active outbreaks during the study period, from March/2021 to August/2022 for outbreak investigation, assessing of the associated risk factors and molecular detection of the FMD virus circulating in this area.

### 2.2. Study Population

Cattle of greater than six months of age and all breeds under different husbandry and management system were the study population for seroprevalence. The study was conducted on cattle that have experienced outbreaks of FMD and manifesting the clinical signs of the disease such as vesicles on the dental pad, tongue, muzzle or, hooves, teat for molecular detection of FMDv and serotyping. Cattle more than six months age groups, breed, sex and production system (intensive, semi-intensive and extensive management systems) were considered.

### 2.3. Study Design

A cross sectional study design was employed from March 2021 to August 2022. Active outbreak of FMD reported and a field investigation was conducted at a particular site of outbreaks within the study area. Animals of different age groups, sex and breed that shows clinical symptoms of the disease, history of infection but having healing lesions and any other asymptomatic cattle in the same farm or grazing were included in sample collection.

### 2.4. Ethics Statement

I am writing to provide confirmation regarding the experimentation on animals conducted as part of our research endeavors here at Debre Tabor University. As per the requisite procedures, I am pleased to confirm that all such experiments have been conducted in compliance with established animal welfare guidelines and have received prior approval from our institute’s ethics committee. “Ethics approval number DTU2534/23” Dr. Getnet Yitayih Alemu’’

### 2.5. Sample Size Determination

The sample size for serological study was calculated according to the formula given by Thrusfield (2005) to determine the minimum sample size required for this study. According to the study conducted at Bench Maji zone, having almost similar agro-ecology with the current study area, the prevalence of foot and mouth disease from the blood samples was found to be 12.08% (Esayas *et al*., 2009). Thus, the minimum sample size required using expected prevalence of 12.08% and absolute precision of 5% was 163.

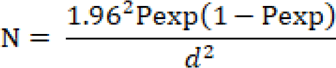

Where, n =sample size, d^2^ =absolute precision, P_exp_ =expected prevalence and Z =statistic for level of confidence = 1.96. Since cluster sampling was used, the sample size was multiplied by the design effect to minimize intra-clustering error term. The design effect was determined using a formula described by Dohoo *et al*. (2003). Thus, the design effect = 1+ ρ (m - 1), where ρ is the intra-cluster correlation coefficient and m is the minimum number of cattle sampled per cluster; accordingly, FAO and OIE (2016) the minimum number of cattle sampled per cluster shouldn’t be less than 26 with Rho of 0.02 in two stage cluster sampling.

Design effect = 1+0.02(26-1) = 1.5

The new sample size (n’) was recalculated by multiplying n = 163 by the design effect (Dohoo *et al*., 2003). Hence, n’ = n x Design effect. Therefore, n’=163×1.5= 245. For the molecular detection overall 29 tissue samples were purposively taken.

### 2.6. Sampling Methods

The study areas (Dedo district and Jimma town) were purposively selected based on presence of active FMD cases, their accessibility and on the abundance of cattle population. Then, cluster sampling was employed to select Kebeles and villages. For seroprevalence study, the villages were considered as clusters (epidemiological units) and simple random sampling method was used to select them. Totally, eight (8) kebeles were selected from the two study areas (four kebeles from each district). The individual cattle from each clusters were selected randomly. The number of cattle sampled from the two-districts were proportional to the total number of cattle population found in the areas. Accordingly, 135 and 110 serum samples were collected from Dedo and Jimma town respectively. From active out break 19 and 10 epithelial tissue and vesicular fluid sample were collected from Dedo and Jimma respectively.

Cattle with evident clinical signs and symptoms of FMD, including a history of oral lesion, history of infection but having healing lesion, and any other asymptomatic cattle (with no evident clinical signs for FMD) in the same farm or grazing with the symptomatic cattle (with evident clinical signs of FMD) were sampled for molecular detection. Data like breed, age, sex, herd composition and size, animal marketing system, the housing system, farming practice were recorded during sample collection. Among husbandry data production systems, history of vaccination, herd size and grazing management were collected to assess their association with FMD sero-positivity. The breed of animal was categorized as cross and local. Age was categorized as 6 months to <3 years of age and >3 years’ age). The production system was categorized as extensive livestock production systems (free grazing), intensive livestock production systems (restricted livestock movement, vehicles, and authorized personnel), and semi-intensive livestock production systems (movement of livestock was less controlled). The vaccination status was recorded as yes or no. Throughout field investigation, data about the disease was gathered by interviewing livestock owners and animal health workers. The data collected was recorded on data collection sheet. Then, animals were clinically examined for presence of FMD lesions on the mouth and feet, and tissue and blood samples were collected.

### 2.7. Sample Collection, Transportation and Processing

About 5ml blood samples were aseptically collected from the jugular vein of apparently healthy animals using plain vacutainer tubes. From clinical cases, at least one gram of tongue epithelial tissues, feet epithelial tissue, and vesicular fluids were collected from non-ruptured or freshly ruptured vesicles, then placed in the bottle with transport medium (an equal amount of buffer saline solution). The samples were labeled with identification numbers and types of tissue. Both samples were transported using an icebox to the Animal Health Institute (Sebeta) for serological and molecular laboratory analysis.

#### 2.7.1. Serological examination

The serum samples were analyzed by ID Screen FMD NSP competition foot and mouth disease virus 3ABC-ELISA (ID Screen^®^ FMD NSP Competition) kit to detect specific antibodies against the non-structural protein of Foot and Mouth Disease virus (FMDV NSP). On the seropositivity/negativity to FMDV antibodies, the outcome variables were categorized based on the results of the 3ABC blocking enzyme-linked immunosorbent assay. Samples were exposed to non-structural FMDV antigen (NSP 3ABC) coated wells on micro titer plates. Then, samples to be tested and the controls were added to the microwells; anti-NSP antibodies if present form an antigen-antibody complex, which masks the virus epitopes. Anti-horseradish peroxidase (HRP) conjugate was added subsequently to the microwells and it fixes the remaining free epitopes forming an antigen-conjugate HRP-complex. The excess conjugate was removed by washing and the substrate solution; 3,3,5,5 - Tetramethylenidine (TMB) was added. The resulting coloration depends on the quantity of specific antibodies present in the samples being tested. In the absence of antibodies, a blue solution appears which becomes yellow after addition of stop solution. In the presence of antibodies, no coloration appears. Within 15 min in the dark, the result was read by micro plate spectrophotometer at 450 nm optical wavelength. The diagnostic relevance of the result was obtained by comparing the OD, which develops in wells containing the samples with the OD from the wells containing the positive control as the ELISA reader read it. To validate the test, the mean OD value of negative control is greater than 0.7 while mean OD for positive control is less than 0.3. Sera with competition percentage ratio for the test sample and negative control (S/N %) less than or equal to 50% were considered positive and S/N% greater than 50% were considered negative. On herd level, sero-prevalence a herd was considered as positive if one or more animals in the herd were seropositive.

#### 2.7.2. Extraction of FMDV RNA

For original samples, an epithelial tissue suspension was prepared by grinding the sample in a sterile pestle and mortar with sterile sand. The epithelial tissue and virus inoculated cell suspensions were centrifuged at 3000 revolutions per minute (rpm) for ten minutes and the supernatant was taken for the molecular work. Total viral RNA was extracted from the digested tissue supernatant, cell suspension supernatant, and sample using the QIAamp® Viral RNA extraction kit (QiagenInc, Hilden, Germany) following the mini Nucleo Spin column protocol according to the manufacturer’s instructions. The extracted RNA was reverse transcribed to detect the presence of FMD viral genetic material using the one-step real-time RT-PCR according to the manufacturer’s instructions. The real-time RT-PCR was used to amplify genome fragments of FMD virus in diagnostic materials including epithelium and the probang samples (Callahan *et al*., 2002).

Briefly, 140 μl of the original epithelial tissues and probang sample suspension was added to 560μl buffer AVL plus carrier RNA in the mirco-centrifuge and vortexed for 15 seconds to mix and then incubated at room temperature (25^0^C) for 10 minutes. The tube was briefly centrifuged to remove drops from the inside of the lid. Then, 560μl of ethanol (96%) was added to the sample and mixed by pulse overtaxing for fifteen seconds followed by brief centrifuging to remove drops from the inside lid. Then, 630μl of the solution were applied to the QIAamp Mini column in a 2ml collection tube and centrifuged at 600g (8000rpm) for 1 minute. The filtrate was discarded, and the column was placed in a fresh 2ml collection tube. Then, 500μl of buffer AW2 were added to the column then centrifuged at 14,000rpm for three minutes and the filtrate was discarded. Then 65μl of buffer AVE was added to the column equilibrated at room temperature for one minute then centrifuged at 6000g (800rpm) for one minute. Finally, RNA was evaluated in a final volume of 60μl and stored at −80°C. Then after, the RNA sample was used for detection and molecular characterization of FMD virus.

#### 2.7.3. The rRT-Polymerase Chain Reaction to detect viral RNA

The diagnostic real-time PCR indicated that the successfully amplified target gave an amplification curve and the cycle threshold (Ct), at which the target amplicon was initially detected above the background fluorescent levels as determined by the embedded software. Analysis of the specimen samples using diagnostic real-time PCR for FMDV genome detection was performed for fifty-eight CPE positive FMDV cell culture isolates. The presence of viral RNA was screened by using real-time RT-PCR method targeting the universal 3D regions of FMD virus using specific primers: 3D forward (5′-ACT GGG TTT TAC AAA CCT GTGA-3’) and 3D reverse (5′-GCG AGT CCT GCC ACG GA −3′); and the TaqMan 3D probe (6-FAM 5’-TCC TTT GCA CGC CGT GGG AC −3’ TAMRA) (Callahan *et al*., 2002).

A master mix assay was used with the final volume per reaction as 20.0µL, plus 5.0µL from each template RNA. Briefly, for each sample PCR reaction mix was prepared by considering a reaction mix per one reaction as follows: 2× reaction mix at 12.5μl, nuclease-free water at 1.5μl, 3D forward primer at 2.0μl, 3D reverse primer at 2.0μl, 3D TaqMan probe at 1.5μl and superscript III RT/Platinum TaqMix at 0.5μl. Asia1 was used as positive control and RNase-free water was used as the negative control. The PCR thermal regime for amplification of VP1 domain was as follows: 30 minutes at 60°C (reverse transcriptase step), ten minutes at 95°C (inactivation reverse transcriptase/activation DNA polymerase), fifteen seconds at 95°C(denaturation), and one minute at 60°C (annealing and elongation), with the two last steps performed for 50 cycles.

Measurement of fluorescence was taken at the end of second step at stage 3. Cycle threshold or crossing point (Ct) for each sample was then determined. The cycle threshold (Ct-value) corresponds to the number of cycles required for a given sample to reach the threshold above which it is considered positive. Using a positive cut-off Ct-value of 32.0 (Shaw *et a*l., 2007), FMDV genome was detected by rRT-PCR and results interpretation was as follows: samples with Ct< 32 are classified as positive, samples with 32 < Ct < 40 is ambiguous and marked for retesting and samples with no Ct (undetected) are classified as negative. The cycle threshold value (Ct value) was fixed automatically from the pre-loaded machine software. FMDV positive samples were further analyzed for virus isolation, serotyping, and VP1 sequencing at World Reference Laboratory (WRL) for FMD, Pirbright, United Kingdom (UK). An OIE-recommended pan-serotype-specific one-step rRT-PCR was used as the reference assay. Each sample was tested in duplicate, using primers and probes reported by Callahan *et al*. (2002) and reagents, parameters, and thermal cycling conditions as per Shaw *et al*. (2007). Positive reactions were defined as those, which gave a detectable Ct value.

### 2.8. Questionnaire Survey

Verbal consent was obtained from the interviewees (respondents) those were willing to give data. Interviews were performed in Afan Oromo and Amharic. The objective of the survey was also explained to them. The survey was conducted by administering the semi-structured questionnaires to 143 household representatives (86 males and 57females) for identifying risk factors. The contents of the questionnaire were mainly focused to collect data about the age, vaccination history, the housing system, grazing and watering management, the animal marketing, movement of animals, raising system and the herd size. Interviewing and sampling were done in collaboration with the two district veterinary offices workers. In addition, photos during sample and data collection in the field as well as during laboratory process are present in.

### 2.9. Data Analysis

The data that were collected after laboratory analysis and questionnaire survey were coded and entered into MS-Excel spreadsheet (Microsoft Corp., Redmond, USA). Then, data from MS-Excel were transported to SPSS 25 statistical software (SPSS Inc., Chicago, IL, USA). The descriptive statistics such as frequencies and furthermore, the statistical associations between the suggested risk factors.

The association between FMD seroprevalence and the relevant risk factors was analyzed with multivariable logistic regression. Univariable analysis was performed and factors with p-value <0.25 were taken forward for multivariable analysis to correct for confounding effects. The strength of the association between outcome and explanatory variables was assessed using the adjusted odds ratios (OR). For all analysis, confidence level (CL) is at 95% and P≤0.05 was set for significance. Collinearity of the variables was analyzed using spearman correlation coefficient. Interaction between the variable taken to multivariable analysis was tested and turned with no interaction. The overall model fitting of the final model was tested using the Hosmer–Lemeshow test.

## 3. RESULTS

### 3.1. Individual Cattle Level Seroprevalence

The total number of cattle sampled in this study were 245 (176 females and 69 males), out of which 55 were seropositive for FMD antibody with an overall seroprevalence of 22.45% (95%, CI: 17.2%-27.7%). According to the result obtained from the present study, higher 25.2% (95% CI=18.6-33.1) and the lower 19.1% (95% CI=12.8-27.4) of sero-prevalence were recorded in Dedo district and Jimma town, respectively. The higher 33.6% (95% CI= (25.8 42.5), and the lower 12% (95% CI: 7.4-18.7) seroprevalences were recorded in Large and small herd size, respectively. The Adult age group were more affected (28.8%; 95% CI: 22.3-35.5) than younger age group (7.2 %; 95% CI: 3.1-15.9). In extensive and intensive management system, seroprevalences of 19.5% (95% CI: 14.8-25.2) and 5% (95% CI=31-68.6) were recorded respectively.

**Table 1:** Overall FMD Seroprevalence and Associated Putative/possible risk factors.

| variables | category | Sample examined | positive sample | Prevalence in % (95% CI) | COR (95% CI) | P-value |
| --- | --- | --- | --- | --- | --- | --- |
| District |  |  |  |  |  |  |
|  | Jimma Town (Ref) | 110 | 21 | 19.1 (12.8-27.4) |  |  |
|  | Dedo | 135 | 34 | 25.2 (18.6-33.1) | 1.43 (0.77-2.64) | 0.257 |
| Kebele |  |  |  |  |  |  |
|  | Mentina (Ref) | 28 | 5 | 17.9 (7.9-35.8) |  |  |
|  | Furustale | 27 | 5 | 18.5 (8.2-36.7) | 1.05 (0.27-4.12) | 0.949 |
|  | Hora Gibe | 27 | 5 | 18.5 (8.2-36.7) | 1.05 (0.27-4.12) | 0.949 |
|  | Ifa Bula | 28 | 6 | 21.4 (10.2-39.5) | 1.26 (0.33-4.71) | 0.737 |
|  | Bilo Adicho | 34 | 9 | 26.5 (14.6-43) | 1.66 (0.48-5.67) | 0.422 |
|  | Odo Hidhata | 33 | 9 | 27.3 (15-33.44) | 1.73 (0.50-5.92) | 0.386 |
|  | Dima Serte | 34 | 9 | 26.5 (14.6-43) | 1.66 (0.48-5.67) | 0.422 |
|  | Keta Qadida | 34 | 7 | 20.6 (10.4-36.8) | 1.19 (0.33-4.27) | 0.787 |
| Agro-ecology |  |  |  |  |  |  |
|  | High-land (Ref) | 177 | 39 | 22 (16.6-28.7) |  |  |
|  | Mid-land | 68 | 16 | 23.5 (15-34.8) | 1.09 (0.56-2.11) | 0.801 |
| Breed |  |  |  |  |  |  |
|  | Cross (Ref) | 46 | 8 | 17.4 (9-30.7) |  |  |
|  | Local | 199 | 47 | 23.6 (18-30) | 1.47 (0.64-3.37) | 0.364 |
| Herd size |  |  |  |  |  |  |
|  | Small (5-20) (Ref) | 126 | 15 | 12 (7.4-18.7) |  |  |
|  | Large (>20) | 119 | 40 | 33.6 (25.8-42.5) | 3.75 (1.94-7.25) | 0.000 |
| Sex |  |  |  |  |  |  |
|  | Male (Ref) | 69 | 13 | 18.8 (11.4-29.6) |  |  |
|  | Female | 176 | 42 | 24 (18.2-30.7) | 1.35 (0.67-2.71) | 0.398 |
| Age |  |  |  |  |  |  |
|  | Young (0.5-3)(Ref) | 69 | 5 | 7.2 (3.1-15.9) |  |  |
|  | Adult (>3) | 176 | 50 | 28.8 (22.3-35.5) | 5.08 (1.93-13.36) | 0.001 |
| New animal marketing |  |  |  |  |  |  |
|  | No (Ref) | 183 | 21 | 11.5 (7.6-16.9) |  |  |
|  | Yes | 62 | 34 | 54.8 (42.5-66.6) | 9.37 (4.77-18.41) | 0.000 |
| Management system |  |  |  |  |  |  |
|  | Intensive (Ref) | 24 | 12 | 5 (31-68.6) |  |  |
|  | Extensive | 221 | 43 | 19.5 (14.8-25.2) | 4.14 (1.74-9.85) | 0.001 |
| Overall |  | 245 | 55 | 22.5 (17.7-28) |  |  |

Ref, Reference; COR, Crude Odds Ratio; CI, Confidence Interval

The univariable logistic regression analysis on individual animal-level risk factors showed that the herd size (P=0.000), age (P=0.001), new animal marketing (P=0.000) and management system (P=0.001) had statistically significant effect on sero-prevalence of FMD (P<0.05). No significant interaction between variables was detected. Accordingly, they were selected for the final model.

**Table 2:** Multivariable Logistic Regression analysis for risk factors association with sero-prevalence of FMD in in animals of Dedo and Jimma Districts.

| Factors | Sample examined | positive sample | Prevalence in % (95% CI) | AOR (95% CI) | P-value |
| --- | --- | --- | --- | --- | --- |
| Herd size |  |  |  |  |  |
| Large (>20) | 119 | 40 | 33.6 (25.8-42.5) | 3.8 (1.8-8.2) | 0.001 |
| Small (5-20) (Ref) | 126 | 15 | 12 (7.4-18.7) |  |  |
| Age |  |  |  |  |  |
| Adult (>3) | 176 | 50 | 28.8 (22.3-35.5) | 3.75 (1.3-11.2) | 0.018 |
| Young (0.5-3)(Ref) | 69 | 5 | 7.2 (3.1-15.9) |  |  |
| New animal marketing |  |  |  |  |  |
| Yes | 62 | 34 | 54.8 (42.5-66.6) | 7.53 (3.6-15.8) | 0.000 |
| No (Ref) | 183 | 21 | 11.5 (7.6-16.9) |  |  |
| Management system |  |  |  |  |  |
| Extensive | 221 | 43 | 19.5 (14.8-25.2) | 5.1 (1.7-15) | 0.004 |
| Intensive (Ref) | 24 | 12 | 5 (31-68.6) |  |  |
| **Overall** | **245** | **55** | **22.5 (17.7-28)** |  |  |

Ref, Reference; AOR, Adjusted Odds Ratio; CI, Confidence Interval

From the final model, herd size (P=0.001), age (P=0.018), new animal introduction (P=0.000) and the management system (P=0.004) estimated to be potential risk factors for FMD sero-positivity. Hence, cattle with large herd size were almost 4-times (OR=3.8) more likely to be sero-positive for FMDV antibody than the smaller herd size. Adult cattle had nearly 4-times (OR=3.75) odds of being sero-positive compared to young. Animals from herds to which a new animals was introduced were in the odds of almost 8-times (OR=7.53) to be seropositive compared to those animals from the herds to which no new animals were added. Cattle reared under extensive management system were 5-times (OR=5.1) more likely seropositive than those raised in intensive.

### 3.2. Questionnaire Survey Analysis

Cattle owners were asked about the condition of their animals; 143 owners who provided their animals for sampling were interviewed. Out of the respondents, 64% replied that their cattle were affected by a disease with a clinical sign of depression, excessive salivation, lameness and sore lesion on tongue, pad, interdigital space and mouth in the last one year. According to the finding of the questionnaire survey, when animals become sick, 81% of the interviewee replied, when get diseased their animals, they consult animal health officials to get the treatment while 19% of the respondents treat their animals based on their own home grown knowledge towards the disease by purchasing drugs either from animal health post or markets. Among the respondents, there was a practice of mixing different herd group during watering or grazing. Most of the respondents (75%) use markets to sale their live animals while 8% sale either directly to nearby neighbor or to intermediaries. The rest of the interviewee (17%) used both selling in the market and nearby village or neighbor.

Out of the respondents, 82% buy live animals from local nearby markets, 13% purchase from nearby village or neighbor. The remaining 5% use both markets and nearby villages for purchasing animals. The majority of the respondents believed that the main source of FMD outbreak was communal grazing and watering points, and unrestricted movement of animals. In considering of the vaccination history of the sampled animals against FMDV, 47% of the respondents replied that their animals have never had vaccination against FMD where as 15% of the interviewee does not exactly differentiate between vaccine and ordinary drug they had taken. However, 38% of the respondents replied they had vaccine against FMD. Most of the respondents had been witnessed FMD outbreaks among their cattle. Of the respondents that witnessed FMD outbreaks 72% of them had one outbreak in a year. The rest experienced two (16%) and three (12%) outbreaks per year. during the outbreak, 80% of the respondents, reported no deaths of cattle showing the clinical symptoms.

### 3.3. Molecular Detection of FMDV

The successfully amplified target gave an amplification curve and the cycle threshold (Ct). The target amplicon was initially detected above the background fluorescent levels determined by the embedded software. The Ct value (cycle threshold or crossing point) corresponds to the number of cycles required for a given sample to reach the threshold above is considered as positive. The real-time RT-PCR (using universal primers and probe of FMDV) was performed on all 29 collected samples from Dedo district (n=19) and Jimma town (n=10). Among 29 tested samples, 27 (93.1%) samples were found positive (17 from Dedo and 10 from Jimma town), having a Ct value ranging from 17.01 to 36.79 and the fluorescence of samples rises above the background fluorescence and were serotype SAT 2.

**Figure 2:**
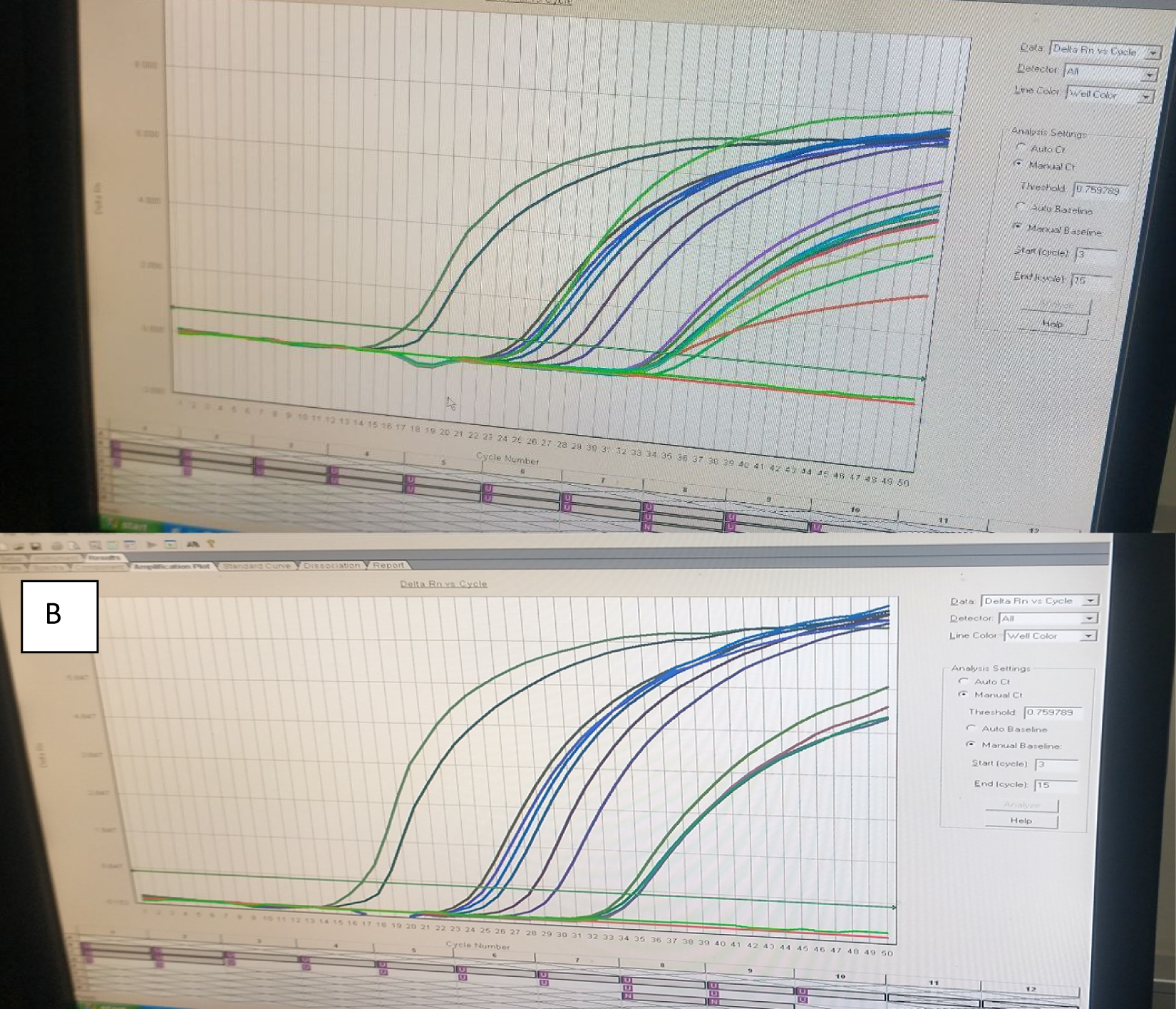
Amplification curve and the cycle threshold (Ct) sample from (A) Dedo district and (B) Jimma town.

**Table 3:** Molecular Detection of FMD Virus and Serotyping from Samples of Outbreak Cases with Ct value.

| No: | Sampling area | Sex | Age | Sample item | Ct value | Identified Serotype |
| --- | --- | --- | --- | --- | --- | --- |
| 1 | Jimma Town | Male | 5 | Epithelial Tissue | 25.13 | SAT2 |
| 2 | Jimma Town | Female | 2 | Epithelial Tissue | 17.00 | SAT2 |
| 3 | Jimma Town | Female | 10 | Epithelial Tissue | 34.90 | SAT2 |
| 4 | Jimma Town | Male | 6 | Epithelial Tissue | 18.49 | SAT2 |
| 5 | Jimma Town | Female | 2 | Epithelial Tissue | 28.22 | SAT2 |
| 6 | Jimma Town | Female | 8 | Epithelial Tissue | 24.76 | SAT2 |
| 7 | Jimma Town | Female | 6 | Epithelial Tissue | 26.44 | SAT2 |
| 8 | Jimma Town | male | 4 | Epithelial Tissue | 30.85 | SAT2 |
| 9 | Jimma Town | female | 7 | Epithelial Tissue | 28.62 | SAT2 |
| 10 | Jimma Town | male | 5 | Epithelial Tissue | 24.45 | SAT2 |
| 11 | Dedo | Male | 4 | Epithelial Tissue | 29.73 | SAT2 |
| 12 | Dedo | Female | 2.5 | Epithelial Tissue | 34.65 | SAT2 |
| 13 | Dedo | Female | 7 | Epithelial Tissue | 34.16 | SAT2 |
| 14 | Dedo | Female | 3 | Epithelial Tissue | 36.33 | SAT2 |
| 15 | Dedo | Female | 6 | Epithelial Tissue | 34.78 | SAT2 |
| 16 | Dedo | Female | 1 | Epithelial Tissue | 25.88 | SAT2 |
| 17 | Dedo | Female | 9 | Epithelial Tissue | 31.37 | SAT2 |
| 18 | Dedo | Female | 5 | Epithelial Tissue | 35.74 | SAT2 |
| 19 | Dedo | Female | 1.5 | Epithelial Tissue | 19.45 | SAT2 |
| 20 | Dedo | Female | 3 | Epithelial Tissue | 36.79 | SAT2 |
| 21 | Dedo | Female | 1 | Epithelial Tissue | 34.21 | SAT2 |
| 22 | Dedo | Male | 5 | Epithelial Tissue | 35.04 | SAT2 |
| 23 | Dedo | Male | 2 | Epithelial Tissue | 34.37 | SAT2 |
| 24 | Dedo | Female | 2 | Epithelial Tissue | 31.50 | SAT2 |
| 25 | Dedo | Female | 5 | Epithelial Tissue | 26.08 | SAT2 |
| 26 | Dedo | Female | 5.5 | Epithelial Tissue | 35.56 | SAT2 |
| 27 | Dedo | Male | 4 | Epithelial Tissue | 29.73 | SAT2 |

## 4. DISCUSIONS

The present study showed that the overall individual cattle level seroprevalence of FMD was 22.5%. This figure is in agreement with the results of 21.4% from Borena and Guji by Sulayeman *et al*., 2018, 24.2% from central Ethiopia by Desissa *et al*., 2014, 24.6% from Borena by Rufael *et al*., 2008 and 26.5% from Kellem Wolega by Mekonen *et al*., 2011. The current finding was higher than the findings of 9% by Abunna *et al*., 2013 from western Ethiopia, 8.1% by Belina *et al*., 2016 from Dire Dawa, 9.5% by Molla *et al*., 2010 from Southern Ethiopia. Also higher than 8.9% by Mesfine *et al*., 2019 South Omo Zone, 10.9% by Megersa *et al*., 2009 East Shewa Zone and 14.4% by Beyene *et al*., 2015 Amhara region. On the other hand, Mekonnen *et al*., 2005, Kibore *et al*., 2013 and Lazarus *et al*., 2012 from Ethiopia, Kenya, and Nigeria with the prevalence of 53.6%, 52.5%, and 72.6%, respectively, reported higher seroprevalence of FMD compared to this study result. These differences in sero-prevalence might have emerged from differences in sampling size, study design, and the presence and absence of extrinsic risk factors like agro-ecology, contact of animals with wildlife, free movement of animals, communal grazing and communal watering (Rufael *et al*., 2008; Abunna *et al*., 2013; Desissa *et al*., 2014; Abdela, 2017; Sulayeman *et al*., 2018).

In this study, herd size based sero-prevalence was determined. Large herd size was more likely to be sero-positive than the smaller herd group. It was also indicated that large herd size (33.6%) had significantly higher sero-positivity than the small herd size (12%). This observation is agreeing with the reports of Gelaye *et al*., 2009 from Benchi Maji zone, Desissa *et al*., 2014 from Kellem Wolega, Sulayeman *et al*., 2018 from central Ethiopia, Molla *et al*., 2010 from South Omo zone and Mesfine *et al*., 2019 from Amhara region. This direct association might be an indication of contagious nature of the disease and mode of transmission, which is attributed to crowding of animals that can facilitate frequency of direct contact and hence enhance the likelihood chances of transmission (Ayelet *et al*., 2012; Gorna *et al*., 2014; Saeed *et al*., 2015; Endris, 2018).

From the multivariable logistic regression, the chance of being sero-positive was higher in the adults (>3 years). The adult cattle had higher sero-prevalence (28.8%) than the younger age group (7.2%). This finding is in line with the findings of Dubie and Negash, 2021, who found a higher prevalence in adult animals than in young animals in a study conducted at the Afar region. Other researchers who documented the same finding were Awel and Dilba., 2021 in Addis Ababa, Megersa *et al*., 2009 in Southern Ethiopia, Sulayeman *et al*., 2018 in central Ethiopia, Gelana, 2016 in Western Oromia, and Abunna *et al*., 2013 in Dire Dawa. These statistically significant prevalence differences between different age groups reported might be due to increased exposure to disease risk factors as an animal’s age increases. Calves < 1 year are protected from the disease due to their passive maternal immunity (Tesfaye *et al*., 2016).

In contrast to previous reports by Rufael *et al*., 2008 in Borena pastoral area, Megersa *et al*., 2009 in Gamo Gofa and Sidama Zones and Ahmed *et al*., 2020 in West Shewa zone Oromia Regional State revealed significantly higher FMD seroprevalence in young as compared with adult and old cattle. On the other hand, Gelaye *et al*., 2009 in the Benchi Maji zone and Belina *et al*., 2016 in the Eastern Showa zone reported no statistically significant difference in the seroprevalence of FMD in different age groups. The variation in reports might be due to variation in sample size allocation among age categories and variation in management system of young animals; separate housing of young animals or keeping young animals apart from adult animals around the homestead (Mekonen *et al*., 2011; Lazarus *et al*., 2012; Beyene *et al*., 2015; Sulayeman *et al*., 2018; Ahmed *et al*., 2020).

The final multivariable logistic regression analysis showed that the likelihood of being FMD seropositive for newly introduced cattle had nearly 8-times (OR=7.53) more chance than home borne and this practice was a potential risk factor for the occurrence of FMD (P=0.000). These results are in agreement with the report of Ayelet *et al*., 2012, FAO, 2012 and Smith *et al*., 2015. For the FMD-endemic countries, an incursion of an exotic FMD virus is likely to result in extensive regional spread because of intense intraregional livestock trade. This might be farmers do not seek veterinary support before mixing newly purchased animal into the herd. In addition, animals are taken to market and brought to home on foot crossing long distances. During this stress time, the animals become susceptible for different infections. This may be explained by the fact that when cattle from different origins come together in one market place and there might be interactions and spread FMDV. Consequently, they come home carrying the virus and play a role in the disease transmission (Ayelet *et al*., 2014; Tadesse *et al*., 2018; Ludi *et al.,* 2019).

Management system had shown significant association with FMD seroprevalence, in that it was higher in animals managed under extensive (19.5%) than intensive (5%) farming systems. Moreover, the odds of animals with FMD were 5-times (OR=5.1) higher in extensively managed cattle than intensively managed. This finding is in agreement with a study conducted by Megersa *et al*., 2009, Gelana, 2016, Ayelet *et al*., 2014 and Molla *et al*., 2010 where free grazing was identified as one of the major risk factors for FMD transmission. This may be attributed to a high level of herd mobility, contact of animals at grazing and watering points, dynamism of herds (frequent additions) and frequent contact with the livestock of neighboring peasant associations. This is also due to the risks associated with local spread leading to contamination or the roads and environment close to known infected premises, and the increased risk of undetected infected premises (Rufael *et al*., 2008; Mekonen *et al*., 2011; Beyene *et al*., 2015; Mesfine *et al*., 2019).

In this study, FMD occurrence, its frequency, husbandry practice, control strategy and the attitude of the farmers in the study area were assessed. It was found that frequent occurrence, free type of livestock grazing, uncontrolled trade movements in and out and sharing pasture with herds of different peasant associations and with wildlife were among the practices seen in their location. According to the results of the questionnaire, communal grazing places and watering points, different animals in close contact and presence of nearby market places and failure to vaccinate animals could be the possible factors that contributed to the high disease transmission and occurrence. This finding aligns with assessments made by various authors, indicating that continual exposure to susceptible animals, the mingling of diverse herds, and the shared use of grazing areas and watering points by a large number of susceptible animals are contributing factors to the occurrence of Foot-and-Mouth Disease (FMD) outbreaks (Sahle *et al*., 2004; Abbas *et al*., 2012; Gelana, 2016; Ludi *et al* 2019; Mesfine *et al*., 2019). In the study areas, there was no official FMD control strategy in place, but the practice of different control and prevention measures were in place before and during the occurrence of the disease. Respondents also mentioned the practice of traditional treatment methods used in the area to dress FMD lesions. The finding of Mesfine *et al*., 2019 from the Amhara region where the traditional wound management system was observed, supported this.

In this work, out of 29 bovine epithelial tissue samples tested by real-time PCR, 27 were found positive for 3D regions of FMDV. The negative findings for two samples may be due to the small amount of viral RNA in the initial samples, which may have caused sample dissemination during lab work. Inline to this work, some researchers documented that the preferred sample for virus detection is the epithelial tissue, and there is higher levels of viral RNA in the epithelial tissues (OIE, 2012; Getachew *et al*., 2017). According to Shaw *et al*., 2007, the epithelial tissue samples from the vesicular lesions could be used as the sample of choice for FMDV detection.

In the current study, the detected FMDV serotype from 27 epithelial tissue samples was serotype SAT 2, which is an indication of predominant serotype circulating in sites of outbreaks from Dedo district and Jimma town, South-West Ethiopia. In contrast to this finding, previous studies showed that serotype O was the most prevalent and dominant serotype causing outbreaks in the different parts of the Ethiopia (Ayelet *et al*., 2009; Negussie *et al*., 2011; Sulayeman *et al*., 2018).

## 5. CONCLUSSION AND RECOMMENDATIONS

The present study was undertaken with the objective of investigation of the sero-prevalence, and associated risk factors and molecular detection of FMDV. The overall sero-prevalence (22.5%) showed that the FMDV infection has been circulating in the study area. The presence of antibodies against FMDV implied exposure to active or previous viral infection. The disease was found to be more prevalent in Dedo district than Jimma town. The study showed herd size, the practice of introducing newly purchased animal and management system were identified as the risk factors or predictors for the occurrence and distribution of FMD infection. From outbreak cases, FMDV serotype SAT-2 was identified using molecular tools. Farmers had knowledge of FMD and the associated clinical signs, but the disease control by vaccination and its coverage reported in this area was low. Traditional and antibiotic treatment was the chief management measure applied in the study areas. As control of FMDV at its source is of common interest; so, national support is very important for the study area. Hence, in reference to the concluding remarks, the following recommendations are pointed out:

➢ Mass vaccination better be practiced in the study areas.
➢ Farmers and livestock owners should be trained on the disease, husbandry practices and how to report FMD outbreaks.
➢ Improved understanding of FMD epidemiology for the farmers can help identify risk-based control measures that can be implemented to reduce disease impact.
➢ Effective monitoring systems should be implemented for timely information about the occurrence, spread, and causative serotype identification.
➢ Further study should be conducted at a wider range on virus isolation, serotyping, and molecular characterization of the FMD virus.

